# Frontopolar cortex is a mediator of network modularity in the primate brain

**DOI:** 10.1101/2019.12.20.882837

**Authors:** Zhemeng Wu, Matthew Ainsworth, Helen Browncross, Andrew H. Bell, Mark J. Buckley

## Abstract

Primate frontopolar cortex (FPC), occupied by area 10, sits atop a functional hierarchy of prefrontal cortical regions yet little is known about its causal role influencing cognition and brain networks. We studied resting-state-functional-connectivity (rsfc) networks in rhesus monkeys with or without FPC; 86 cortical regions showed significant differences in inter-area rsfc measures in lesioned animals versus controls. K-means clustering showed these regions were organized into two distinct networks in lesioned animals, whereas the same areas clustered into four networks in control animals. These networks extended within and beyond prefrontal cortex. These results suggest that FPC is involved in mediating cortical networks in the primate brain, both within and beyond prefrontal cortex. Even after 40 months only partial recovery of lesion-induced reduced modularity had occurred. We therefore suggest that FPC might help implement long-standing diverse neural network dynamics.

## Introduction

Frontopolar cortex (FPC) is occupied by area 10, a cytoarchitectonic area exclusive to the human and non-human primate brain. Studies using non-invasive neuroimaging methods have previously linked human FPC activities with a bewilderingly diverse range of higher-order cognitive functions, to the extent that it has proved impossible to reach consensus from neuroimaging studies on a fundamental role of FPC. In human neuroimaging studies, for example, FPC has been found to be associated with maintaining and integrating subgoals to fulfil the final behaviour outcomes ^1–4^, explorative behaviour ^5,6^, cognitive branching ^7^, multi-tasking ^8^, future planning ^9^, prospective memory ^10–12^ and memory awareness ^13,14^ amongst many others. Clearly, whatever contributions to cognition are supported by FPC, they involve interactions with other posterior brain regions.

Whilst all these cognitive operations must require inter-area coordination, some human studies have explicitly revealed some FPC interactions. For example, Sakai and Passingham (2006) demonstrated that FPC interacts differently with different posterior cortical regions depending on the intended cognitive operation held in working memory ^15^. In a similar vein, in hypothesizing that FPC encodes the evidence in favour of switching to alternative courses of action, Boorman et al. (2009) found that FPC entered into a distinct pattern of functional connectivity with the parietal cortex immediately before a behavioural switch occurred, leading to the proposal that FPC engages the parietal cortex to implement the switch ^6^.

Whether such observations of interactions reflect causal influences on posterior regions cannot be determined from correlational neuroimaging studies as to reveal causality one needs to study interventions. However, naturally occurring selective FPC lesions interventions are extremely rare. Human neuropsychology has traditionally proceeded by considering many patients with varied and overlapping lesions and assigning patients to groups if their lesions encroach significantly on target areas such as FPC irrespective of selectivity. That FPC may have a role in optimizing attentional allocation and controlling or switching attention between alternatives (including in one theory between externally available stimuli and self-generated thoughts) especially in the context of novel, ambiguous, and open-ended situations, or during multi-tasking and cognitive branching, are some of the key insights to have arisen from human neuropsychological investigations of PFC ^16–19^. It is hard to draw firm conclusions for patients with such extensive lesions. One way to address this is take large numbers of patients with non-selective brain lesions and apply voxel-based lesion analyses to see which lesioned voxels are most commonly associated with their behavioural deficits. Using this approach Volle et al. (2011) showed that lesioned voxels in FPC were associated with time-based prospective memory (a task which may be interpreted as one that requires monitoring of the relative value of performing one of two tasks at any given time) ^20^. There exist only extremely rare case studies of individual patients with lesions restricted to within FPC; indeed, we know of only one ^21^ and that adds little significant insight to the issue (other than reported marked indecisiveness and lack of normal response inhibition) as perhaps the most relevant behavioural tests were not administered.

Therefore, we turn to animal models (in non-human primates (NHP) as rodents do not have area 10 of FPC) wherein both highly circumscribed lesions and electrophysiology may be targeted specifically to NHP FPC. Only two published studies have targeted neuronal recordings of FPC and these indicate that while FPC makes little contribution to on-going decision making it appears to be engaged in evaluating decision outcomes prior to feedback ^22,23^, contrary to other prefrontal regions involved in goal-directed behaviour ^23^. Until very recently there were no circumscribed FPC lesion studies in NHPs but three recent studies now indicate that FPC lesions in macaques produce a pattern of spared, impaired, and enhanced behaviour which together may be interpreted as evidence that FPC mediates exploration and rapid learning about the relative value of alternatives of a broad kind ^24–26^. Relatedly, one study administered reversible silencing of a sub-region of macaque PFC, rather than acute lesions of all of FPC as in the above studies, and found that FPC inactivation in this way impaired metacognitive evaluation of non-experienced events ^27^. Moreover, a recent review on the function of FPC, taking into account the NHP and human literature, proposes a functional model of FPC in which FPC contributes to disengaging cognitive control from the current task and redistributing resources to other potential goals for ‘exploration’; while the posterior parts of the prefrontal cortex are involved in staying on the focus on the current task to facilitate ‘exploitation’ ^28^. Setting the optimum balance between ‘explorative’ and ‘exploitative’ tendencies, especially in dynamic volatile environments, is likely to be a key process common across many different cognitive tasks and this is likely to necessitate a close functional connections and interactions between FPC and multiple other prefrontal and posterior brain networks. Anatomical connections between FPC and other brain regions are broadly similar across human and NHPs ^29,30^. However, compared to other prefrontal cortical areas, FPC has a unique pattern of reciprocal connections to other cortical areas in that while its connections to prefrontal cortex are extensive, its connections to posterior regions are not and appear to be exclusive to multimodal regions (i.e. prominent reciprocal projections: *via* external capsule to multimodal regions in superior temporal sulcus and superior temporal gyrus region, *via* uncinate fasciculus to select regions in the anterior and medial temporal lobe including perirhinal cortex, and *via* cingulate fasciculus to anterior and posterior cingulate and retrosplenial cortices ^31^) suggestive that FPC my sit atop a hierarchy of prefrontal regions in the primate brain. Accordingly, we hypothesize that FPC is causally involved in mediating far-reaching networks of brain regions within and beyond prefrontal cortex. This study investigates this using circumscribed FPC lesions and network cluster analyses of whole brain functional connectivity analyses. Resting state functional connectivity (rsfc) examines the correlations in spontaneous brain activities in absence of external stimuli or task demands and offers a tool for understanding the functional organization of whole brain networks. Seven adult macaque monkeys received bilateral lesions to the entire FPC; three of them had MRI scans 2 months post-lesion and four of them had scans 40 months post-lesion ^24^. We first compared extra-FPC rsfc in 13 control animals (i.e. with intact FPC) with the same measures in the FPC lesioned animals scanned 2 months post-lesion to investigate how presence versus absence of FPC affects widespread brain networks (using cluster analyses to distinguish networks). We ran the same analyses with the 40 months post-FPC lesion group to investigate how robust the observed disturbances in FPC-mediated brain networks were across years.

## Results

We collected resting state fMRI data and structural scans under anaesthesia from a total of 20 animals. These included 13 control animals with intact FPC, 3 FPC-lesioned animals scanned 2-month post-lesion and 4 FPC-lesioned animals scanned 40-month post lesion. All seven lesioned animals received complete bilateral aspiration lesions of the FPC (Fig.1). Resting state functional connectivity (rsfc) was calculated between 260 cortical regions by pairwise correlation from the mean BOLD time series.

**Figure 1.**
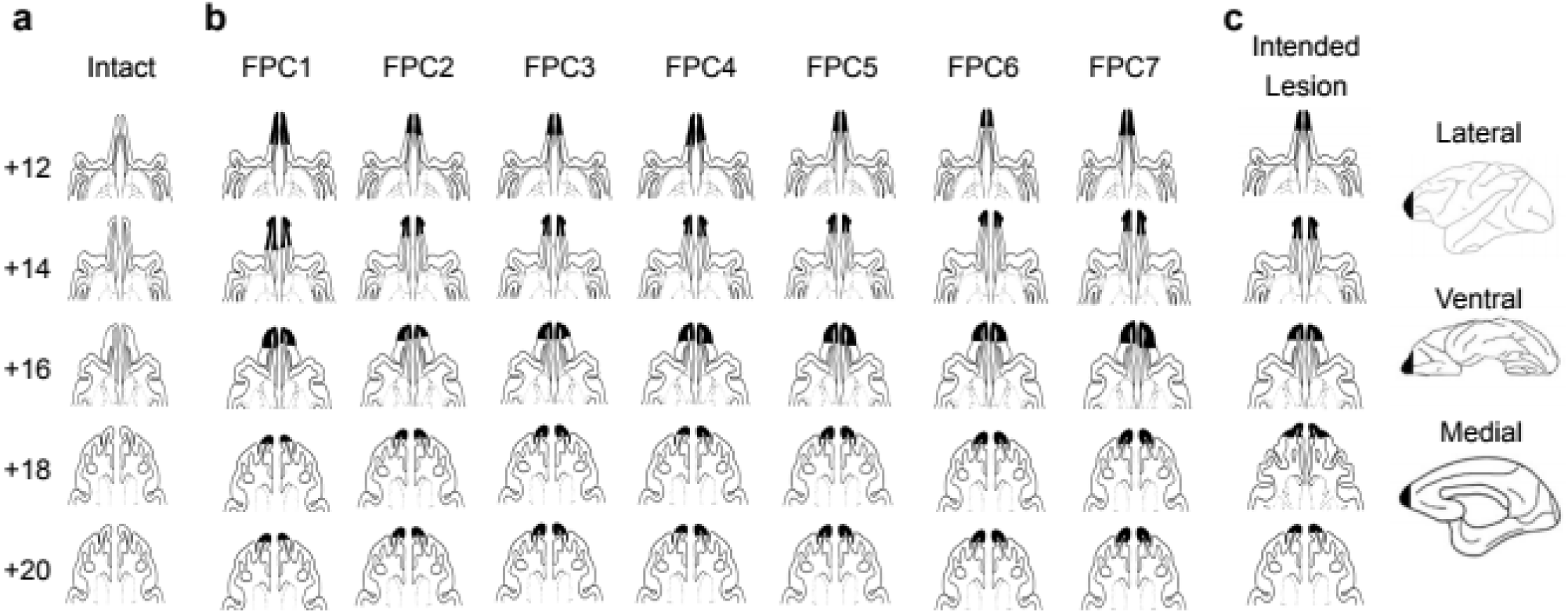
Intended and actual extent of the bilateral FPC lesion. Drawings of horizontal sections of macaque prefrontal cortex showing the intact FPC (**a**), and the actual lesion extent from all seven FPC lesioned animals (**b**, FPC 1-4: 40-month post-FPC lesion group; FPC 5-7: 2-month post-FPC lesion). The intended extent of the lesion is shown on the same horizontal sections and summarised in drawings of the lateral, ventral, and medial surfaces (**c**). Numbers represent the approximate distance in millimeters above the interaural plane.

### Changes of functional connectivity across the brain after circumscribed lesions to FPC

To investigate the changes in functional connectivity after lesions to FPC, we constructed matrices of the pairwise correlations between all 260 cortical regions. Matrices of mean functional connectivity in unoperated control animals versus in 2-month post-FPC-lesioned animals are shown in Fig. 2a and Fig. 2b respectively.

**Figure 2.**
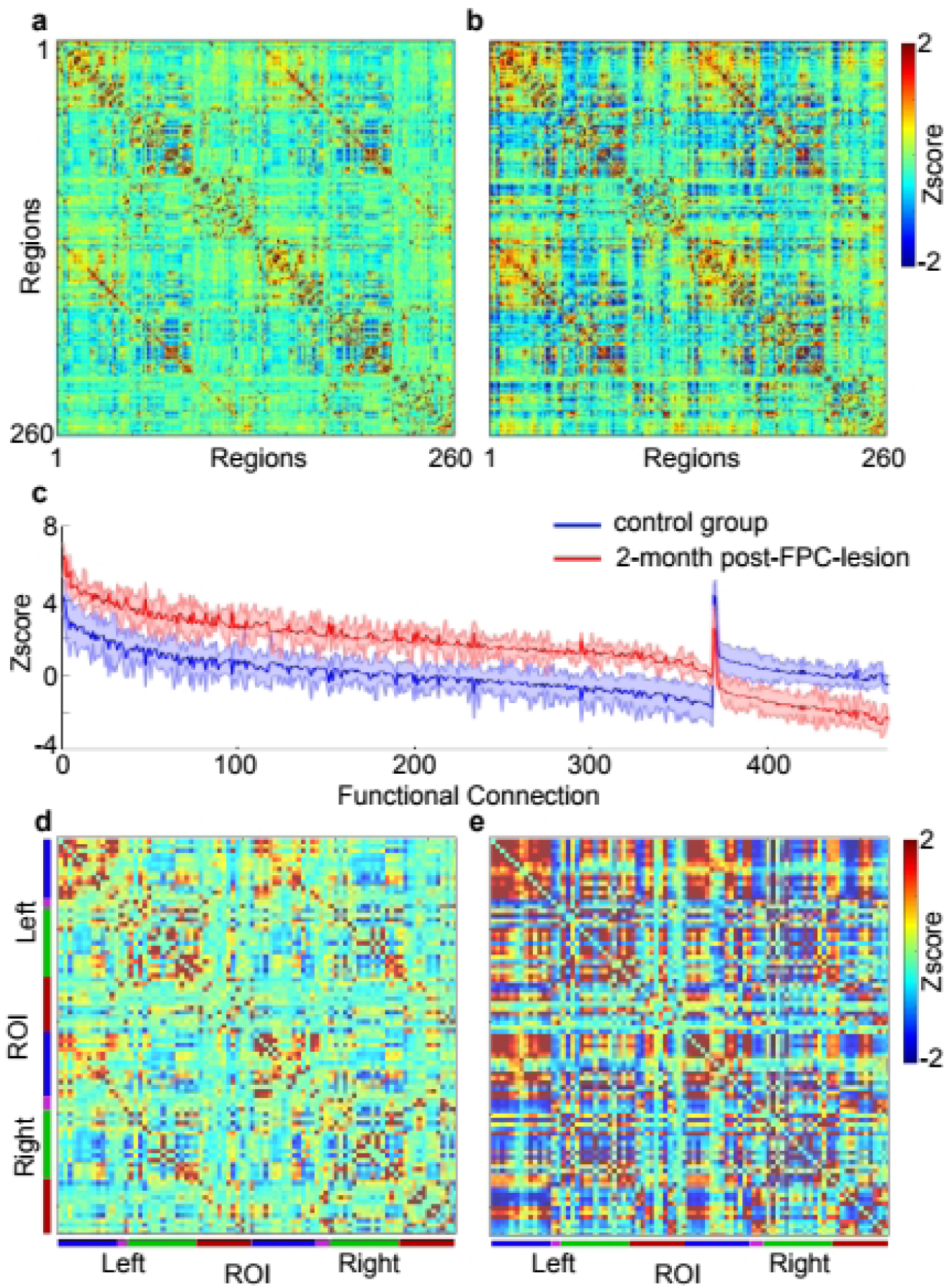
The impact of lesions to Frontopolar cortex on resting state functional connectivity. Matrices of functional connectivity between 260 cortical regions for control group (**a**) and 2-month post-FPC lesion group (**b**). A total of 469 functional connections (linking 86 cortical regions) were revealed by a nonparametric permutation test to be significantly different between the two animal groups (two tailed test, p<0.05, see methods for details of the test). The majority of connections were enhanced in the 2-month post-FPC lesion group (red) when compared with control group (blue, **c**). Matrices of functional connectivity between the 86 identified regions reveal marked differences between the control group (**d**) and 2-month post-FPC lesion group (**e**). Colour bars (left and below matrices) indicate ROIs within the frontal (blue), occipital (pink), parietal (green), and temporal (red) lobes respectively.

To probe significant differences in functional connectivity between the two animal groups, a standard assumption-free nonparametric permutation test was applied to these rsfc data (see methods) ^32^. In this test the difference in connection strength between the lesion and control group was compared to a surrogate distribution of the maximal/minimal difference observed after random re-labelling of the lesion and control groups (repeated 10,000 times). Testing in this way revealed 469 functional connections which were significantly different between control animals and the 2-month post-FPC-lesioned animals (two-tailed permutation test, *p* < 0.05). Of the 469 functional connections deemed to have significantly changed, the majority were found to have enhanced functional connectivity following FPC lesion: 369 connections increased while 100 connections decreased in strength (Fig. 2c). These significant connections linked a subset of 86 cortical regions-of-interest (ROIs), distributed both within and beyond prefrontal cortex (see Table 1 for full list and MRI coordinates of selected ROIs). To further explore changes in connectivity following lesion of FPC we therefore considered only connections between these 86 ROIs (Fig. 2d and 2e functional connectivity matrices).

### Multidimensional scaling reveals a striking lesion effect of FPC

To aid in the visualization of the significant changes in functional connectivity between control animals and 2-month post-FPC-lesioned animals, we applied a multidimensional scaling (MDS) analysis was applied to the pairwise correlations between these 86 identified ROIs in both groups. In this analysis we projected all connections into a 2-dimensional space, where ROIs with similar overall patterns of functional connectivity across the overall matrix are plotted with greater proximity to one-another (Fig. 3 a-b). We then used K-means clustering (with the variance to be explained fixed at 80%) to define clusters within the 2-dimensional space for both the control and the 2-month post FPC-lesioned animals’ scaled data (Fig. 3 a-c).

**Figure 3.**
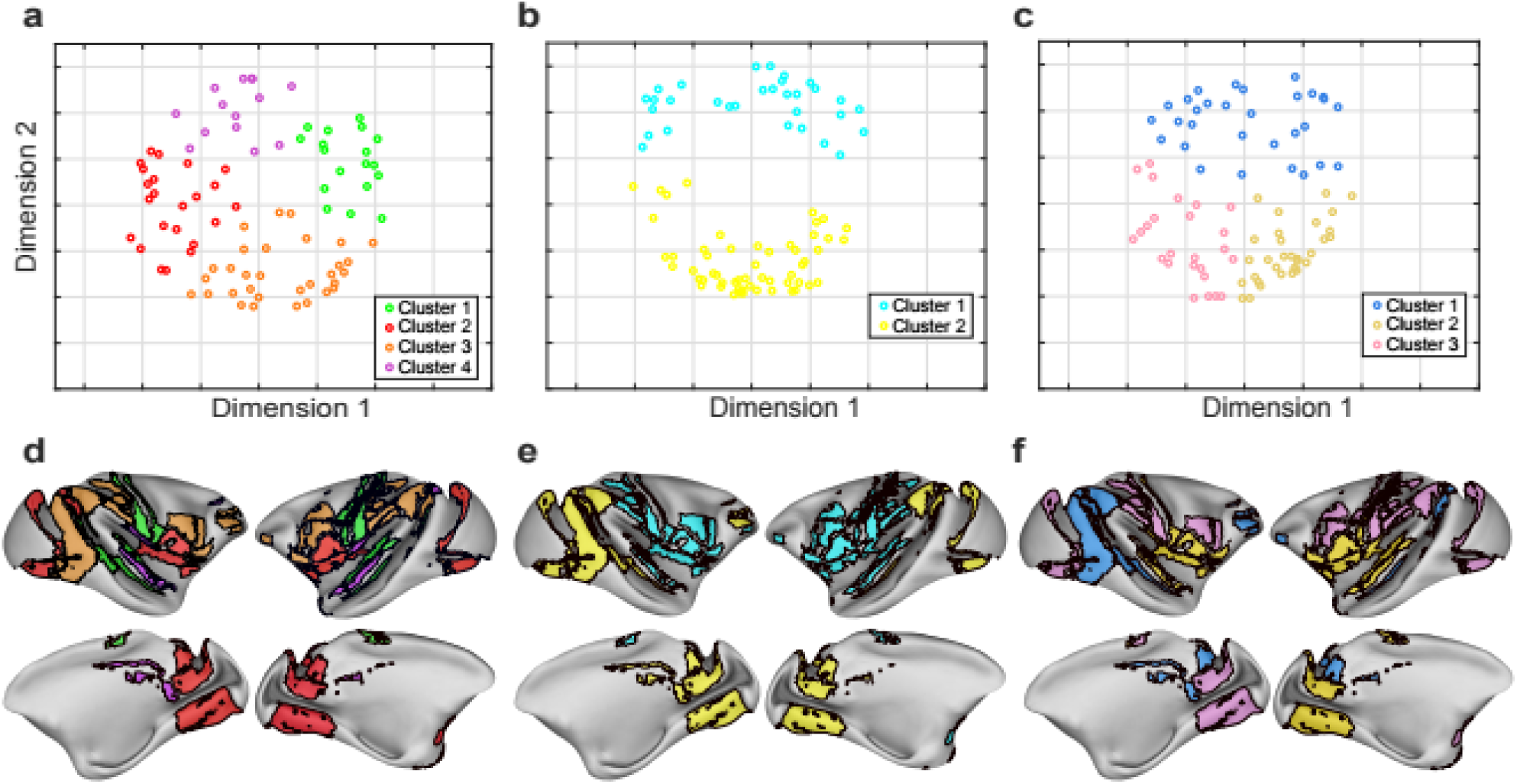
Multidimensional scaling reveals reduced network modularity in the absence of Frontopolar cortex. Multidimensional scaling and cluster analysis of the functional connectivity between the 86 previously identified ROIs for animals in the control group (**a**), 2-month post-FPC lesion group (**b**), and 40-month post-FPC lesion group (**c**). Clustering of ROIs was defined by K-means clustering (into sufficient clusters to explain 80% of total variance). Each ROI is represented with a single dot, with cluster class denoted by colour. The distance between ROIs corresponds to the similarity/dissimilarity in the patterns of functional connectivity of each ROI respectively. Differences in the cluster classes of all ROIs in the control (**d**), 2-month post-FPC lesion (**e**), and 40-month post-FPC lesion group (**f**) was visualised on the surface of the macaque brain, adopting the same colour conventions as in a-c.

In control animals this projection revealed the existence of four clusters (Fig. 3a) whereas only two distinct clusters were evident in the projection of 2-month post-FPC lesion animals (Fig. 3b). Moreover, visualization of the 2-dimensional space of all ROIs in the FPC-lesioned animals revealed a large spatial separation between the two clusters (Fig. 3b). In addition, these clusters showed physical separation in the brain when the anatomical locations of the regions were considered. Specifically, there was a distinct anterior versus posterior clustering of brain regions (Fig 3e). The first cluster included frontal regions, areas of insula cortex, somatosensory cortex, premotor and motor cortex, anterior parietal cortex and regions of the anterior temporal lobe; while the second cluster predominantly included posterior cortical regions including posterior parietal cortex, cingulate cortex, retrosplenial cortex, occipital cortex and posterior temporal lobe regions. In contrast, in the control group, the four clusters show less obvious spatial separation on the 2D plot (Fig. 3a) and less obvious gross anatomical separation between networks (Fig. 3d).

### Differences in cluster-based analyses between 2-month versus 40-month post-FPC lesion groups

To explore how robust are our observed FPC-lesion mediated changes in rsfc clusters (mindful of the possibility of recovery/plasticity in the brain after some brain insults), we compared these results to corresponding rsfc and MDS projections from the 40-month post-FPC-lesion group.

We observed three clusters in the 40-month post-PFC lesion group (Fig. 3c). The spatial (i.e. Fig 3c versus Fig3a/Fig3b) and anatomical (i.e. Fig 3f versus Fig3d/Fig3e) separations appeared to fall between the control group and the 2-month post-FPC lesion group. This suggests some but not complete restoration of functional organization after 40 months compared to control animals.

To better illustrate the similarities and differences in cluster organization between the three groups of different animals (control, 2-month post FPC lesion, 40-month post FPC lesion), we made a 3D plot to link the three 2-dimensional projections for the 86 ROIs across groups (Fig. 4). Fig. 4 shows that the four clusters observed in control animals appear to collapse into two distinct clusters (shown in blue and yellow) in 2-month post FPC-lesion animals, with dramatic changes in the position of individual ROIs. The three clusters, observed in the 40-month post-FPC-lesion group, appeared more integrated than for the two clusters in the 2-month post-FPC-lesion group. However, there were limited changes in the position of individual ROIs between these two group, particularly when compared with the more robust differences observed between controls and 2-month post-FPC-lesion group. This provides evidence that the 40-month period, a relatively long time in terms of brain plasticity and recovery of function, did not result in anything more apparent than a relatively small and partial recovery of network structure.

**Figure 4.**
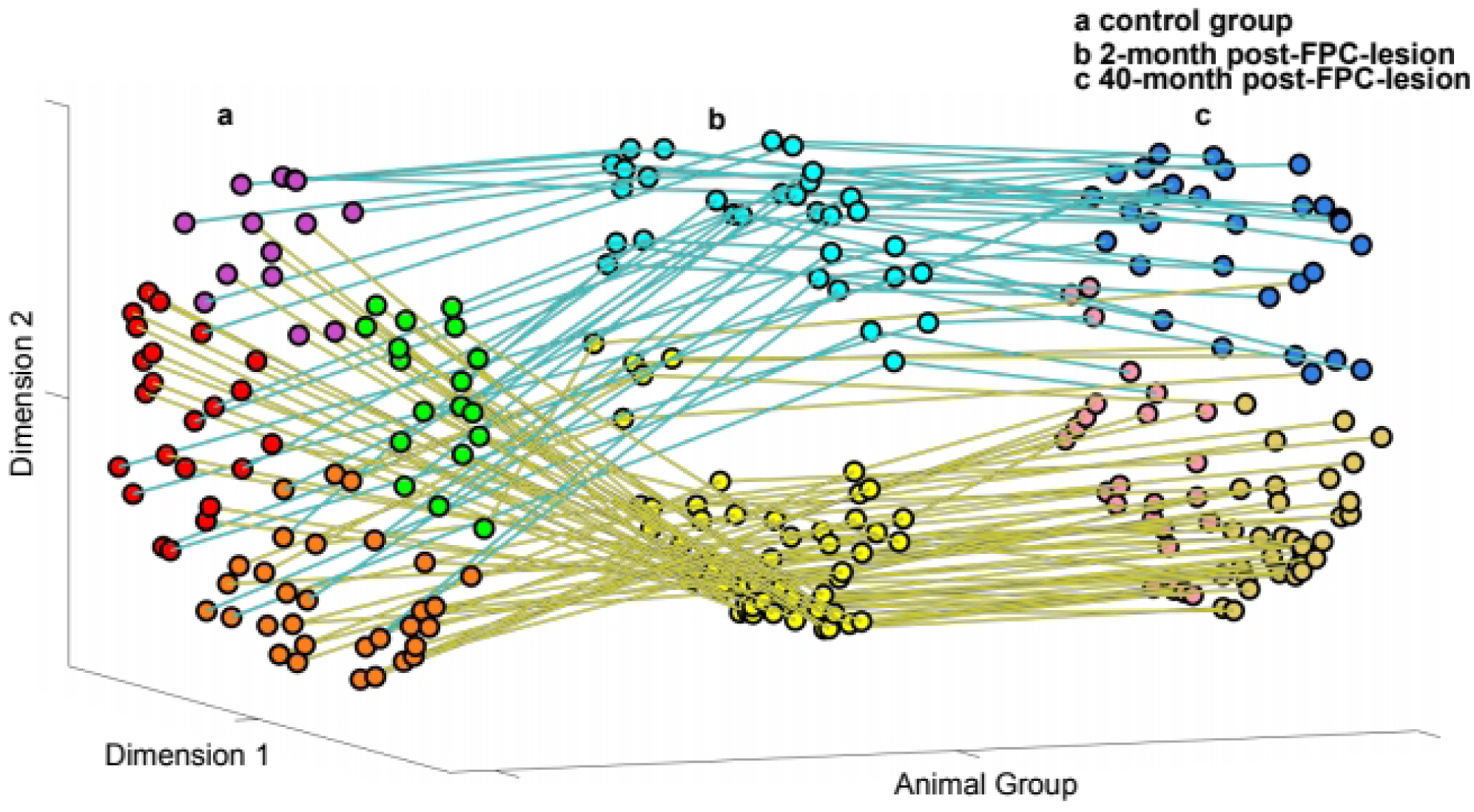
Visualisation of changes in the relationship between ROIs explored by multidimensional scaling. Multi-dimensional scaling plots of the relationship between all 86 ROIs in the control group (**a**), 2-month post-FPC-lesion group (**b**), and 40-month post-FPC-lesion group (**c**). Changes in the location occupied by each ROI are visualised by linking across lesion groups. ROI scatter plots follow the conventions in Figure 3a-c. Line linking ROIs between lesion groups are coloured based on whether the ROI was in cluster 1 (blue) or cluster 2 (yellow) for the 2-month post-FPC-lesion group.

### Predictors of changed rsfc patterns across the brain in absence of FPC

We next sought to enquire whether the specific patterns of observed changes of rsfc induced by FPC lesions might be predicted by brain organization observed in the control animals with intact FPC.

Our first hypothesis is that the magnitude of rsfc between FPC and other ROIs in control animals predicts (i.e., with a positive correlation) the extent of repositioning of ROIs in the 2D projection (and therefore changes in the pattern of connectivity between ROIs) in the FPC lesioned animals. To test this hypothesis, for each of the 86 ROI’s, we computed the Euclidean distance (in 2D space) between the position occupied by that region in control animals and the position occupied by that region in 2-month post-FPC-lesioned animals. Correlation of the Euclidean distance between these two groups, with the absolute connection strength between each ROI and FPC in control animals revealed a significant positive correlation (Pearson’s *r* = 0.233, *p* = 0.031, Fig. 5a). This suggests that the stronger rsfc of one brain region to FPC in control animals, the greater the separation between that regions position in 2-dimensional space in control animals and FPC-lesioned animals. In other words, FPC-induced lesion effects on functional connectivity can predict how brain networks may adapt in the absence of FPC.

**Figure 5.**
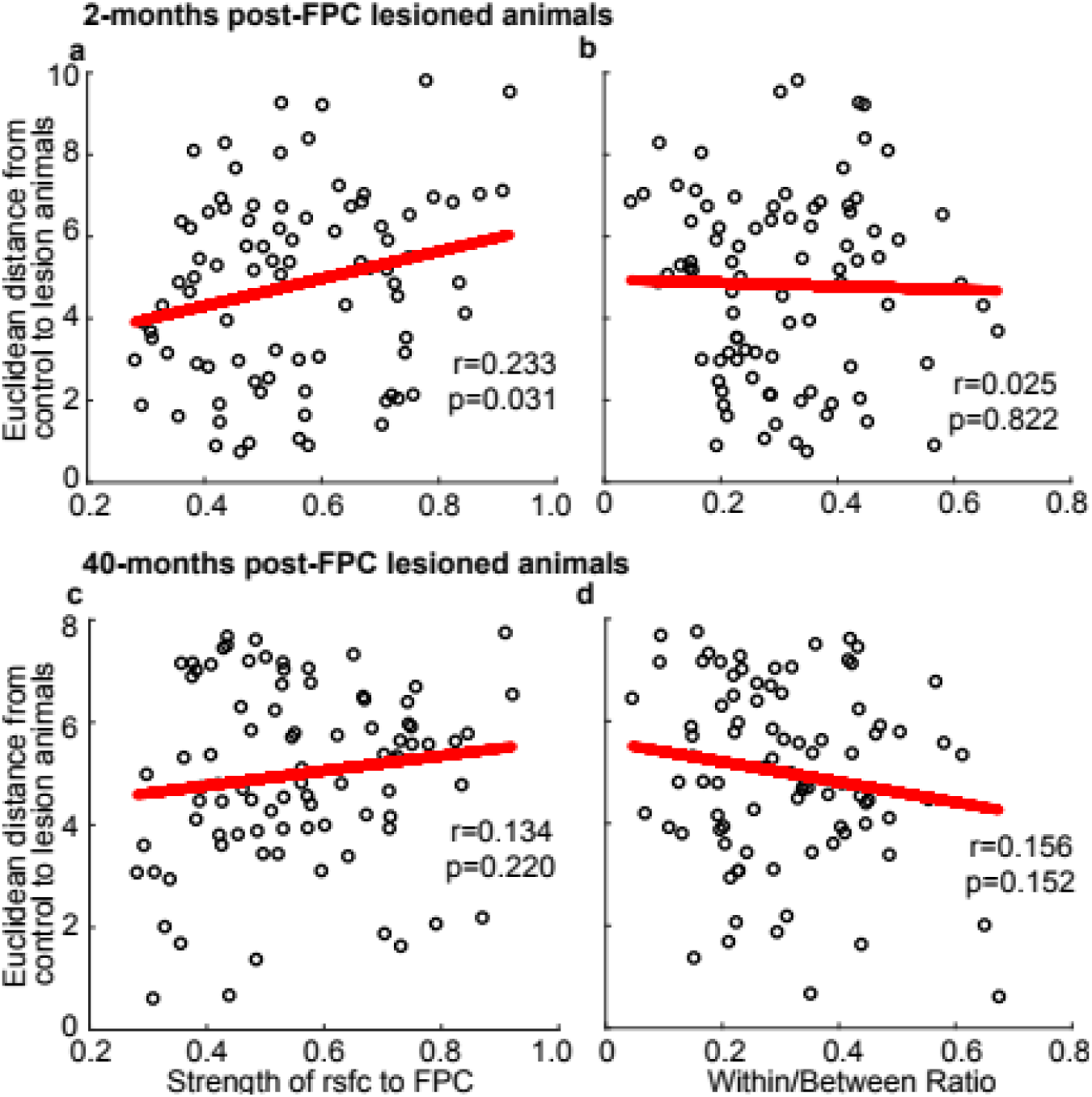
Remodelling of ROI connectivity after lesion of frontopolar cortex is predicted by ROI to FPC connectivity in control animals. Scatter plots showing the relationship between the strength of an ROIs connection to FPC in control animals and the difference between an ROIs position in 2-D space (calculated as Euclidean distance) in control animals vs. in 2-month post-FPC-lesion animals (**a**) and the relationship between the within/between ratio of an ROI in control animals with the Euclidean distance between control animals vs. in 2-month post-FPC-lesion animals (**b**). The same relationships, FPC connection strength vs. Euclidean distance (**c**) and within/between ratio vs. Euclidean distance (**d**) are shown for ROIs between control animals and 40-month post-FPC-lesion animals. An ROIs within/between ratio is defined as the average distance to within cluster ROIs divided by average of distance to ROIs in other clusters). Correlation coefficient (Pearson’s r) and the significance of the correlation shown for each test.

Our second hypothesis is that in the absence of FPC, connectivity between ROIs can be predicted according to the relationship between ROIs in control animals. In this scenario the separation between an ROIs position in 2D space in control and FPC lesioned animals would be dependent on its relative location within the four clusters observed in control animals, with ROIs more loosely or peripherally related to the relevant cluster demonstrating the greatest change in position (and therefore connectivity) after FPC lesions. Therefore, we investigated whether *centrality* versus *peripherality* of a ROI within the four clusters observed in control animals might predict the change in rsfc between a ROI and other areas after FPC lesions. To calculate the relative location of each ROI in the 2-dimensional scale in control animals, we defined an index, called the within/between ratio. This ratio was calculated as the average of distances (in the 2-dimensionally scaled data) between each ROI and other ROIs within their local cluster divided by average of distances between the ROI and other ROIs in other clusters; hence the within/between ratio was larger for ROIs situated towards the margins of its own cluster and smaller if the ROIs were located more centrally in its own cluster. We did this for each of the 86 ROIs in control animals compared to the same ROIs in the 2-month post-FPC lesioned animals. Correlation of the within/between ratio with the Euclidean distance between the control and 2-month post-FPC lesioned animals was not significant (Pearson’s *r* = 0.025, *p* = 0.822, Fig. 5B), therefore our second hypothesis was not supported.

We also tested these two hypotheses in the 40-month post-FPC-lesioned animals, as compared to controls, and found that neither of the correlations was significant (strength of rsfc to FPC: Pearson’s *r* = 0.134, *p* = 0.220; within/between ratio: Pearson’s *r* = 0.156, *p* = 0.152, Fig. 5c and 5d). This suggests that the network state that exists in the 40-month post-FPC lesion animals and that likely includes any plasticity-mediated recovery of function occurring over a few years is no longer predictable from the aforementioned parameters of the control animals network state. This further suggests that even after several years, plasticity such that it exists is insufficient to return the network to a nearly normal state. Hence the effects of FPC lesions on extended brain networks may be robust for several years (and although our data cannot speak to longer periods than 40 months, 40 months appears to be a long time in terms of brain plasticity and recovery of function, especially in light of our own previous studies of functional recovery after frontal lesions in macaques ^33^, so it is not unreasonable to predict that FPC-lesion mediated changes to brain network structure may indeed extend to decades or even to the lifetime).

## Discussion

Circumscribed bilateral lesions to the macaque FPC produced a distinctive change in rsfc across multiple brain networks extending both within and beyond prefrontal cortex. Analyses of the patterns of changes suggest FPC lesions reduce the number of distinct networks as compared to the intact animal. In control animals our cluster-based analyses of rsfc revealed the following four networks: (i) a primary auditory cortex, somatosensory, occipital, superior and middle temporal cortical network (cluster 1); (ii) a gustatory cortex, piriform cortex, ventral premotor cortex, insular, occipital, posterior parietal and middle temporal cortical network (cluster 2); (iii) a ventral prefrontal, premotor, motor, somatosensory, occipital, anterior areas of posterior parietal and perirhinal cortical network (cluster 3); and (iv) a posterior prefrontal, somatosensory, superior temporal sulcal, cingulate, and retrosplenial cortical network (cluster 4). Notably each of these clusters extended over multiple cortical lobes. This indicates that animals with an intact FPC can maintain several distinct and complex cortical networks. By contrast, in the absence of FPC, the ROIs collapsed into two brain networks: a frontal-temporal-anterior parietal network and a cingulate-occipital-posterior parietal network, with each demonstrating disparate resting state connectivity patterns according to our cluster analyses. The frontal-temporal-anterior parietal network occupied more anterior parts of the brain, which included prefrontal cortex, premotor and motor cortex^34,35^, gustatory cortex, piriform cortex, somatosensory cortex, insular regions, anterior regions in parietal cortex, anterior regions in temporal cortex and perirhinal cortex. This brain network is typically associated with executive control ^36^, somatosensory and olfactory perception ^37,38^, motion and body movement ^39^ and mnemonic processing ^40,41^. The cingulate-occipital-posterior parietal network occupied more posterior parts of the brain, which included posterior cingulate cortex, retrosplenial cortex, posterior parietal cortex, occipital cortex, medial superior temporal areas and middle temporal areas. The regions involved in these networks are typically associated with emotional ^42^, visual ^43,44^, and spatial processing ^45^.

The relationship between functional connectivity and anatomical connectivity is not precise (directly connected areas need not oscillate in synchrony, and it is not necessarily the case that areas without direct anatomical connections cannot exhibit synchrony as posterior areas without direct connections to FPC may be strongly connected to other posterior areas and ‘recruited’ into a FPC-mediated networks indirectly). Nonetheless, anatomical tracing studies (see below) and functional connectivity studies in macaques ^46,47^ do indicate that many brain regions are in fact both anatomically and functionally connected to FPC. Petrides and Pandya (2007), Markov et al. (2014) used anatomical tracing to identify cortical and subcortical efferent and afferent connections to and from FPC in macaques ^31,48^. Despite the different techniques and different species used, a broadly similar range of connections were found in these tracing studies. FPC has dense interconnections with other prefrontal areas (e.g. area 8Ad, 9, 11, 12, 13, 14 and 32); and robust cortical connections with temporal lobe regions, including insula, rostral superior temporal gyrus and superior temporal gyrus regions (e.g. area Ts, TAa and TPO) *via* the external capsule; and anterior and medial temporal lobe (e.g. perirhinal cortex) *via* the uncinate fasciculus; and anterior and posterior cingulate and retrosplenial cortex ^31,48^ *via* the cingulate fasciculus. Moreover, Markov et al. (2014) identified a much wider range of posterior connections with area 10, including connections with parietal and occipital cortex, in which connections were observed with lateral, medial and posterior inter-parietal and with V2 ^48^, although these connections have not been observed in other studies and in some cases the projections are very light. In summary, these evidence of cortico-cortical connections to/from FPC in tracing studies showed a wide range of cortical regions are involved in the cortical networks related to FPC, and most of them have been detected in the frontal-temporal-anterior parietal and cingulate-occipital-posterior parietal network in our rsfc study.

As reviewed in the introduction, the three studies that have investigated macaques with bilateral and circumscribed lesions to FPC have demonstrated that FPC is necessary for considering the relative values of (unchosen) alternatives of a broad kind. This can lead to deficits in some contexts (e.g., specific impairments in the one-trial learning phase of both concurrent objects-in-scenes learning and successive single problem learning wherein animals chose between novel alternative stimuli; and impairments in learning to choose between novel alternative abstract rules ^24^) yet can also lead to enhancements in other contexts (e.g., macaques with FPC lesions showed enhancements in a Wisconsin Card Sort Test analog involving conflict between two well learnt rules in terms of being less distractible than control animals by to intentional distraction between trials and hence better ongoing performance on the focused-upon task ^25^). Together these studies showed that FPC is necessary for disengaging executive control from the current task/goal/stimuli and redistributing it to explore any other potential tasks/goals/stimuli ^25,28^. It is understandable then that the kinds of behavioural changes induced by FPC lesions are both broad in scope, yet also specific to certain kinds of situations such as those where exploration versus exploitation of chosen versus unchosen alternatives is required. Accordingly, the effects of FPC lesions cannot be simply understood as a loss of function of the specific area removed; to understand FPC function one has to consider how FPC causally interacts with a broad range of posterior areas.

The current study contributed to this goal by demonstrating how brains with an intact and functioning FPC influence network cohesion are able to maintain more posterior networks compared to brains without a functioning FPC. We propose that FPC may be considered a mediator of modularity. We show that in addition to changes in behaviour following FPC lesions ^24–26^, we also observe changes to rsfc. But we do not know if the robust changes in rsfc patterns themselves had any influence upon (long-term) behaviour, or whether changed behaviour influenced long-term rsfc (or both).

Speaking to the latter possibility in broad terms it has been shown that such effective connectivity can at least predict cognitive flexibility in humans ^49^. Speaking to the former, resting state networks have themselves been linked to memory consolidation processes that may also be active in the resting state ^50^ and indeed, patients exhibiting episodic memory loss who had lesions to more polar than dorsolateral prefrontal areas have been observed to have greater deficits in remembering than knowing ^51^. Given we have shown a clear causal influence of FPC lesions on posterior network organization, the next set of studies should investigate the influences of FPC on posterior network organization during behaviour itself by conducting rsfc in the awake state. Causal influences of dynamic network states on behaviour may be determined by selective intervention in the dynamic network states (e.g. *via* microstimulation to drive or disrupt oscillations in target areas) in the presence and absence of FPC.

As the control and FPC-lesioned animals scanned after 2 months versus after 40 months post-lesion demonstrated distinct patterns of brain network organizations, we investigated what, if any, factors inherent to network organization on controls might influence network organization in FPC-lesioned animals. Our supported hypothesis, that brain regions with the greatest change in their connectivity pattern following an FPC lesion (measured as change in position in 2-D space) were those that exhibited stronger effective connectivity to FPC in control animals. This was true for 2-month post-FPC lesion but no longer true in the 40-month post-PFC lesioned animals. These findings imply that FPC is important not only in influencing select key areas that it robustly interacts with but is also important for shaping wider posterior brain networks involving those key areas. After 40 months without the influence of FPC the brain’s network organization has evolved, this is unsurprising, but the fact that the post-lesion network structure has not returned to normality after a few years only indicates how FPC plays a critical long-standing role in brain functional organization in primates.

Animals with an intact FPC can maintain more distinct sets of brain networks than animals without FPC. Moreover, the functional connections of regions robustly interacting with FPC are predictive of the changed network structure within and beyond prefrontal cortex in absence of FPC. The predictive changes last at least 2 months and even after 40 months the network structure has not returned to normality. Future work should investigate how these FPC-related brain networks in awake behaving animals influence essential cognitive processes, such as learning, memory and decision making by combining neuroimaging and electrophysiology with reversible targeted interventions during behaviour.

## Materials and Methods

### Animals

Resting state fMRI and structural scans were collected under anaesthesia from seven macaque monkeys (Macaca mulatta) after bilateral lesions of FPC (area 10) (Fig. 1). Monkeys were scanned either 2 months post-operatively (three female monkeys with an average age and weight of 4.4 years and 7.8 kg at time of scanning) or 40 months post operatively (four female monkeys with an average age and weight of 8 years and 6.7 kg at time of scanning). In addition, we collected resting-state fMRI data from a control group of thirteen monkeys (one female and twelve males, with an average age and weight of 4.5 years and 7.6 kg at time of scanning). All animals were socially housed in enriched environments with a 12hr light/dark cycle and had *ad libitum* water access. Prior to participation in this study all animals received basic training on an identical set of behavioural tasks reported elsewhere ^24^. All animal surgery, anaesthesia and experimental procedures were carried out in accordance with the guidelines of the UK Animals (Scientific Procedures) Act of 1986, licensed by the UK Home Office, and approved by Oxford’s Committee on Animal Care and Ethical Review.

### Surgery

Bilateral lesions of FPC (macaque area 10) was carried out by aspiration of cortical tissue under sterile conditions. The caudal limit of the lesion on the dorsolateral surface was 2 mm posterior to the rostral tip of the principal sulcus. On the orbital and medial cortical surfaces, the lesion extended caudally to match the same anterior-posterior extent of the lesion on the dorsolateral surface. Whilst removing tissue care was taken to preserve sub-cortical white-matter where possible. See Boschin et al., 2015 for further details of surgical procedures ^24^.

### Histology

At the conclusion of the experiments, the FPC-lesioned animals were deeply anesthetized and then perfused through the heart with saline, followed by formol-saline solution. Their brains were blocked in the stereotaxic plane, removed from the skull, put in sucrose-formalin solution until the block sank, and subsequently cut in 50-μm horizontal sections on a freezing micro-tome. Every m horizontal sections on a freezing micro-tome. Every fifth or tenth section was retained and stained with cresyl violet. The histological slides so prepared and mounted were subsequently scanned at high resolution from which drawings of the actual extent of the lesion were prepared alongside matched drawings of the intended extent (Fig. 1). Such drawings of four of the FPC lesioned animals (those in the 40-month post-FPC lesion group) have already been presented previously ^24^ and are represented again here, now with accompanying drawings of the additional three FPC lesioned animals (those in the 2-month post-FPC lesion group).

### Anaesthesia and MRI data acquisition

The fMRI data in the two animal groups were collected under sevoflurane anaesthesia. Anaesthesia was induced with intramuscular injection of ketamine (10 mg/kg), xylazine (0.125-0.25 mg/kg) and midazolam (0.1 mg/kg) two hours prior to the start of scanning, which allowed ketamine to leave in the monkey’s bodies before data acquisition. After induction, anaesthesia was maintained using sevoflurane. Commonly used volatile anesthetics including sevoflurane have been shown to modulate functional connectivity in the brain in a dose dependent manner ^52^. Therefore, to preserve connectivity and minimize variation in the data EPI data was collected during light anaesthesia, with a targeted sevoflurane concentration of 2.3% across the entire project. There was no significant difference between the sevoflurane concentrations while data was collected for control animals (mean = 2.0%), FPC-lesioned animals with 2-month scanning interval (mean = 2.4%), and FPC-lesioned animals with 40-month scanning interval (mean = 2.4%, *F*_(2,17)_ = 1.225, *p* > 0.05, *η*^*2*^ = .126). Physiological parameters (e.g. heart rate and blood pressure) were used to assess the depth of anaesthesia throughout the scanning sessions (For further details of anesthesia maintenance and monitoring see Mars et al., 2011 & Mitchell et al., 2016) ^53,54^.

fMRI scans were conducted using a horizontal 3T scanner using a custom-made 4-channel phased array coil (H. Kolster, MRI Coil Laboratory, Laboratory voor Neuron Psychofysiologie, KU Leuven). High-resolution structural scans (0.5 mm isotropic voxels) were acquired using a T1-weighted high-resolution magnetization-prepared rapid-acquisition gradient echo (MPRAGE) sequence (128 slices, TR=2500ms; TE=4.01ms). Whole-brain resting-state EPI fMRI data were collected at a 2×2×2mm resolution (36 axial slices, TR=2000 s, TE=19 ms), with an approximate duration of 54 minutes for each animal (1600 volumes).

### MRI data pre-processing and calculation of functional connectivity

All fMRI data pre-processing and analysis was conducted using a combination of MATLAB (The MathWorks Inc.), SPM8 (Statistical Parametric Mapping; www.fil.ion.ucl.ac.uk/spm), FSL (fMRI of the Brain (FMRIB) Software Library; http://fsl.fmrib.ox.ac.uk/fsl/fslwiki/), Caret (Computerized Anatomical Reconstruction Toolkit ^55^) and aa software (automatic analysis ^56^; www.automaticanalysis.org). High resolution structural images for each animal were aligned to a standard space (F99 Rhesus macaque template -in the space of the atlas of Saleem and Logothetis (2012) ^57^ using affine and nonlinear transformations). Images were subsequently segmented into grey matter, white matter and cerebrospinal fluid (CSF) masks ^58^. Resting-state fMRI data were aligned to standard space through a two-stage process and then spatially-smoothed with a 3mm Gaussian kernel (full-width half maximum). Grey-matter masks were defined on a monkey by monkey basis as voxels with grey-matter probability > 0.5 within each animal.

Physiological noise covariates were calculated by extracting up to 6 principal components (sufficient to explain 99% of the variance) from the BOLD time-series for white-matter and CSF tissue masks ^59^. An additional vascular covariate was defined as the mean time-course within a mask encompassing the superior sagittal sinus. A motion covariate was calculated from the EPI data (prior to smoothing and registration) and was defined as the time-course of the average displacement over the expected brain volume. These confound time-series and their first derivatives were then regressed from each grey-matter time-series, and the mean time-series was calculated for each cortical region within the derived from the CC atlas (the LV-FOA-PHT composite atlas) for macaques ^55^. Finally, functional connectivity was estimated by calculating pairwise correlation coefficients (Pearson’s r) for each pair of CC areas (260 areas, 130 in each hemisphere) and transformed using Fisher’s r-to-z transform. For further details of the pre-processing of structural images & resting state data see Ainsworth et al., 2018 & Mitchell et al., 2016) ^33,54^.

### Nonparametric connectivity analysis

To calculate significant changes in functional connectivity between control and 2-month post-FPC-lesioned animals, we tested the mean difference between the two groups using a standard assumption-free nonparametric permutation-based approach ^60^. The test involved a comparison of the observed difference against a reference distribution of differences, under the null hypothesis of no significant change in the functional connectivity between the two animal groups. For each connection, the reference distribution was obtained by performing 10,000 permutations on the animal labels to randomly assign them to two groups. On each loop of the permutation the mean functional connectivity between these randomly assigned groups was calculated and only the minimal and maximal difference was stored. This resulted in both a minimal and maximal matrix (each 260 by 260). The upper and lower thresholds were defined as the 97.5th percentile of the maximal matrix and the 2.5th percentile of the minimal matrix, respectively. Any observed mean difference between the two animal groups greater than the upper threshold or smaller than the lower threshold was deemed significant at the 0.05 level (*p* < 0.05, two-tailed). By selecting the maximal/ minimal value from the permutation distribution, this two-sided nonparametric permutation-based test was sensitive to both positive and negative changes in connectivity and therefore controlled for global type I errors associated with multiple comparisons.

### Multi-dimensional scaling analysis

Further analysis was carried out all possible connections between 86 ROIs revealed by the previous non-parametric analysis. To visual changes in connectivity between these regions multi-dimensional scaling analysis was applied to the connectivity matrices from all 20 animals in the study. For each experimental group (control, 2-month post-FPC-lesion, and 40-month post-FPC-lesion) 3 dissimilarity matrices was calculated from the mean connectivity matrix from each group (normally distribution). Classical multi-dimensional scaling was then carried to generate an x and y position for each ROI in 2-D space. Clustering of ROIs based on these 2-D coordinates was then calculated by K-means clustering. The number of clusters for each group was found using the elbow method, and set to explain 80% of total variance.

To provide a more precise explanation of what drives changes in an ROIs connectivity in the absence of FPC further correlation analysis was conducted. In this analysis the Euclidean distance between an ROIs position in space in the control group and both post lesion groups were compared with two rules; firstly the strength of an ROIs connection to FPC in control animals and secondly an ROIs within/between ratio. An ROIs within/between ratio is defined as the average distance to within cluster ROIs divided by average of distance to ROIs in other clusters). The relationship between these two rules (FPC-connectivity and within/between ratio) and the change in an ROIs position was tested by calculation of a correlation coefficient (Pearson’s r) and the significance of the correlation tested by a standard general linear model.

## Acknowledgements

This work was supported by Wellcome Trust Strategic Award Grant (Ref: WT101092MA) and MRC Project Grant (Ref: MR/K005480/1). We thank G. Daubney for histological support. We thank the team of expert animal technicians and veterinary staff and anaesthetists for their very high standards of animal care and husbandry throughout.

## References

1. Badre, D. & Wagner, A. D. Selection, Integration, and Conflict Monitoring. Neuron 41, 473–487 (2004).

2. Bunge, S. A., Helskog, E. H. & Wendelken, C. Left, but not right, rostrolateral prefrontal cortex meets a stringent test of the relational integration hypothesis. Neuroimage 46, 338–342 (2009).

3. Christoff, K. et al. Rostrolateral prefrontal cortex involvement in relational integration during reasoning. Neuroimage 14, 1136–1149 (2001).

4. Christoff, K., Ream, J. M., Geddes, L. P. T. & Gabrieli, J. D. E. Evaluating Self-Generated Information: Anterior Prefrontal Contributions to Human Cognition. Behav. Neurosci 117, 1161–1168 (2003).

5. Daw, N. D., O’Doherty, J. P., Dayan, P., Seymour, B. & Dolan, R. J. Cortical substrates for exploratory decisions in humans. Nature 441, 876–879 (2006).

6. Boorman, E. D., Behrens, T. E. J., Woolrich, M. W. & Rushworth, M. F. S. How Green Is the Grass on the Other Side? Frontopolar Cortex and the Evidence in Favor of Alternative Courses of Action. Neuron 62, 733–743 (2009).

7. Koechlin, E., Basso, G., Pietrini, P., Panzer, S. & Grafman, J. The role of the anterior prefrontal cortex in human cognition. Nature 399, 148–151 (1999).

8. Burgess, P. W. Strategy application disorder: The role of the frontal lobes in human multitasking. Psychol. Res 63, 279–288 (2000).

9. Burgess, P. W., Gonen-Yaacovi, G. & Volle, E. Functional neuroimaging studies of prospective memory: What have we learnt so far? Neuropsychologia 49, 2246–2257 (2011).

10. Koechlin, E., Corrado, G., Pietrini, P. & Grafman, J. Dissociating the role of the medial and lateral anterior prefrontal cortex in human planning. Proc. Natl. Acad. Sci. U. S. A 97, 7651–7656 (2000).

11. Okuda, J. et al. Thinking of the future and past: The roles of the frontal pole and the medial temporal lobes. Neuroimage 19, 1369–1380 (2003).

12. Okuda, J. et al. Differential involvement of regions of rostral prefrontal cortex (Brodmann area 10) in time- and event-based prospective memory. Int. J. Psychophysiol 64, 233–246 (2007).

13. Ryals, A. J., Wang, J. X., Polnaszek, K. L. & Voss, J. L. Hippocampal contribution to implicit configuration memory expressed via eye movements during scene exploration. Hippocampus 25, 1028–1041 (2015).

14. Yokoyama, O. et al. Right frontopolar cortex activity correlates with reliability of retrospective rating of confidence in short-term recognition memory performance. Neurosci. Res 68, 199–206 (2010).

15. Sakai, K. & Passingham, R. E. Prefrontal set activity predicts rule-specific neural processing during subsequent cognitive performance. J. Neurosci 26, 1211–1218 (2006).

16. Burgess, P. W. Rostral prefrontal cortex: What neuroimaging can learn from human neuropsychology. Mind and the frontal lobes: Cognition, behavior, and brain imaging (New York: Oxford University Press, 2012). doi: 10.1093/acprof

17. Pollmann, S. et al. Selective visual dimension weighting deficit after left lateral frontopolar lesions. J. Cogn. Neurosci 19, 365–375 (2007).

18. Roca, M. et al. The role of Area 10 (BA10) in human multitasking and in social cognition: A lesion study. Neuropsychologia 49, 3525–3531 (2011).

19. Dreher, J. C., Koechlin, E., Tierney, M. & Grafman, J. Damage to the fronto-polar cortex is associated with impaired multitasking. PLoS One 3, (2008).

20. Volle, E., Gonen-Yaacovi, G., de Lacy Costello, A., Gilbert, S. J. & Burgess, P. W. The role of rostral prefrontal cortex in prospective memory: A voxel-based lesion study. Neuropsychologia 49, 2185–2198 (2011).

21. Hoffmann, M. & Bar-On, R. Isolated frontopolar cortex lesion: A case study. Cogn. Behav. Neurol 25, 50–56 (2012).

22. Tsujimoto, S., Genovesio, A. & Wise, S. P. Evaluating self-generated decisions in frontal pole cortex of monkeys. Nat. Neurosci 13, 120–126 (2010).

23. Tsujimoto, S., Genovesio, A. & Wise, S. P. Neuronal activity during a cued strategy task: Comparison of dorsolateral, orbital, and polar prefrontal cortex. J. Neurosci 32, 11017–11031 (2012).

24. Boschin, E. A., Piekema, C. & Buckley, M. J. Essential functions of primate frontopolar cortex in cognition. Proc. Natl. Acad. Sci. U. S. A 112, E1020–E1027 (2015).

25. Mansouri, F. A., Buckley, M. J., Mahboubi, M. & Tanaka, K. Behavioral consequences of selective damage to frontal pole and posterior cingulate cortices. Proc. Natl. Acad. Sci. U. S. A 112, E3940–E3949 (2015).

26. Mansouri, F. A., Buckley, M. J., Fehring, D. J. & Tanaka, K. The Role of Primate Prefrontal Cortex in Bias and Shift Between Visual Dimensions. Cereb. Cortex 1–15 (2019). doi: 10.1093/cercor/bhz072

27. Miyamoto, K., Setsuie, R., Osada, T. & Miyashita, Y. Reversible Silencing of the Frontopolar Cortex Selectively Impairs Metacognitive Judgment on Non-experience in Primates. Neuron 97, 980-989.e6 (2018).

28. Mansouri, F. A., Koechlin, E., Rosa, M. G. P. & Buckley, M. J. Managing competing goals - A key role for the frontopolar cortex. Nat. Rev. Neurosci 18, 645–657 (2017).

29. Barbm, H. & Pandya, D. N. Architecture and Intrinsic Connections of the Prefrontal Cortex in the Rhesus Monkey 375, (1989).

30. Sallet, J. et al. The organization of dorsal frontal cortex in humans and macaques. J. Neurosci 33, 12255–12274 (2013).

31. Petrides, M. & Pandya, D. N. Efferent Association Pathways from the Rostral Prefrontal Cortex in the Macaque Monkey 27, 11573–11586 (2007).

32. Nichols, T. & Holmes, A. Nonparametric Permutation Tests for Functional Neuroimaging. Hum. Brain Funct. Second Ed 25, 887–910 (2003).

33. Ainsworth, M. et al. Functional reorganisation and recovery following cortical lesions: A preliminary study in macaque monkeys. Neuropsychologia 119, 382–391 (2018).

34. Morecraft, R. J. & van Hoesen, G. W. Frontal granular cortex input to the cingulate (M3), supplementary (M2) and primary (M1) motor cortices in the rhesus monkey. J. Comp. Neurol 337, 669–689 (1993).

35. Morecraft, R. J., Stilwell-Morecraft, K. S., Ge, J., Cipolloni, P. B. & Pandya, D. N. Cytoarchitecture and cortical connections of the anterior insula and adjacent frontal motor fields in the rhesus monkey. Brain Res. Bull 119, 52–72 (2015).

36. Cochin, S. Observation and execution of movement: Similarities demonstrated by quantified electroencephalography. Eur. J. Neurosci 11, 1839–1842 (1999).

37. Stettler, D. D. & Axel, R. Representations of Odor in the Piriform Cortex. Neuron 63, 854–864 (2009).

38. De Araujo, I. E. & Simon, S. A. The gustatory cortex and multisensory integration. Int. J. Obes 33, S34–S43 (2009).

39. Kiyoshi Kurata. Corticocortical inputs to the dorsal and ventral aspects of the premotor cortex of macaque monkeys. Neurosci. Res 12, 263–280 (1991).

40. Suzuki, W. A. & Naya, Y. The Perirhinal Cortex. Annu. Rev. Neurosci 37, 39–53 (2014).

41. Brown, M. W. & Aggleton, J. P. Recognition memory: What are the roles of the perirhinal cortex and hippocampus? Nat. Rev. Neurosci 2, 51–61 (2001).

42. Vogt, B. A. Pain and emotion interactions in subregions of the cingulate gyrus. Nat. Rev. Neurosci 6, 533–544 (2005).

43. Boussaoud, D., Desimone, R. & Ungerleider, L. G. Subcortical connections of visual areas MST and FST in macaques. Vis. Neurosci 9, 291–302 (1992).

44. Kaas, J. H. & Collins, C. E. The primate visual system system (2004).

45. Lewis, J. W. & Van Essen, D. C. Corticocortical connections of visual, sensorimotor, and multimodal processing areas in the parietal lobe of the macaque monkey. J. Comp. Neurol 428, 112–137 (2000).

46. Neubert, F. X., Mars, R. B., Thomas, A. G., Sallet, J. & Rushworth, M. F. S. Comparison of Human Ventral Frontal Cortex Areas for Cognitive Control and Language with Areas in Monkey Frontal Cortex. Neuron 81, 700–713 (2014).

47. Neubert, F. X., Mars, R. B., Sallet, J. & Rushworth, M. F. S. Connectivity reveals relationship of brain areas for reward-guided learning and decision making in human and monkey frontal cortex. Proc. Natl. Acad. Sci. U. S. A 112, E2695–E2704 (2015).

48. Markov, N. T. et al. A weighted and directed interareal connectivity matrix for macaque cerebral cortex. Cereb. Cortex 24, 17–36 (2014).

49. Chén, O. Y. et al. Resting-state brain information flow predicts cognitive flexibility in humans. Sci. Rep 9, 1–16 (2019).

50. Albert, N. B., Robertson, E. M., Mehta, P. & Chris Miall, R. Resting state networks and memory consolidation. Commun. Integr. Biol 2, 530–532 (2009).

51. Wheeler, M. A. & Stuss, D. T. Remembering and knowing in patients with frontal lobe injuries. Cortex 39, 827–846 (2003).

52. Peltier, S. J. et al. Functional connectivity changes with concentration of sevoflurane anesthesia. Neuroreport 16, 285–288 (2005).

53. Mars, R. B. et al. Diffusion-weighted imaging tractography-based parcellation of the human parietal cortex and comparison with human and macaque resting-state functional connectivity. J. Neurosci 31, 4087–4100 (2011).

54. Mitchell, D. J. et al. A Putative Multiple-Demand System in the Macaque Brain. J. Neurosci 36, 8574–8585 (2016).

55. Van Essen, D. C., Glasser, M. F., Dierker, D. L. & Harwell, J. Cortical parcellations of the macaque monkey analyzed on surface-based atlases. Cereb. Cortex 22, 2227–2240 (2012).

56. Cusack, R. et al. Automatic analysis (aa): Efficient neuroimaging workflows and parallel processing using Matlab and XML. Front. Neuroinform 8, 1–13 (2015).

57. Saleem, K. S. & Logothetis, N. A combined MRI and histology atlas of the rhesus monkey brain in stereotaxic coordinates (2012).

58. McLaren, D. G. et al. A population-average MRI-based atlas collection of the rhesus macaque. Neuroimage 45, 52–59 (2009).

59. Behzadi Y, Restom K, Liau J & Liu TT. A component based noise correction method (CompCor) for BOLD and perfusion based fMRI. Neuroimage 37, 90–101 (2007).

60. Nichols, T. & Holmes, A., Nonparametric Permutation Tests for Functional Neuroimaging. Hum. Brain Funct Second Ed. 25, 887–910 (2003).

